# When Velocity Becomes Position: Proprioceptive Contributions to State Estimation

**DOI:** 10.64898/2026.01.09.698577

**Authors:** Erik Skjoldan Mortensen, Mark Schram Christensen

## Abstract

In this study, we investigate the relative contributions of positional- and velocity-based proprioceptive feedback to state estimation when visual feedback of the hand is unavailable. We performed a virtual reality experiment in which healthy human participants (N = 22) completed two tasks: a continuous target-tracking task and a sequential reaching task. In both cases, visual feedback of the hand position was withheld, except for a brief moment, during which an offset to the left or right was sometimes applied. By analysing the movement path following the offset visual feedback and comparing it to the non-offset condition, we can infer the extent to which the brain relies on rate-of-change information compared to absolute positional feedback for estimating the hand position. We compare and contrast our empirical data with simulated movements from a computational model based on Bayesian sensory integration under varied assumptions of the reliance on positional and velocity-based feedback. Our human and simulated data together indicate a substantial reliance on velocity-based information, as movement offsets induced by biased visual feedback decay slowly during both target-tracking and sequential reaching tasks. The tendency for eventual convergence between offset and non-offset trials demonstrates that absolute positional information is incorporated, albeit slowly.

## 1 Introduction

The brain must estimate the position and movement of the body to guide action. When vision of the hand is unavailable, this estimate depends in part on proprioceptive feedback. Muscle spindle afferents provide information related to both muscle length and its rate of change, raising the possibility that movement-related signals contribute to estimates of limb position.

Here, we investigate how movement-related proprioceptive information contributes to handstate estimation. If position estimates are updated partly from movement-related signals, then a visually induced position offset may persist after visual feedback of the hand is removed, because subsequent estimates continue to build on the previously estimated state. We test this prediction in a continuous target-tracking task and a sequential reaching task, and compare participants’ movements with simulations of Bayesian sensory integration in which the assumed precision of position- and velocity-based feedback is varied.

## 2 Background

### 2.1 Proprioceptive feedback

Proprioception refers to the sense of body position and movement, which arises from a collection of peripheral mechanoreceptors (Tuthill and Azim 2018). Central among these are the type Ia and type II muscle spindle afferents, which are sensitive to stretch of the parent muscle (Proske and Gandevia 2012). Muscle/tendon vibration is one of the classical experimental techniques used to investigate how these sensors contribute to the perceived pose and movement of the body. Vibration of a muscle/tendon increases the firing rate of both of these types of receptors, thereby biasing the perceived position and movement of the vibrated limb, corresponding to stretching of the vibrated muscle (Goodwin et al. 1972; Eklund 1972; Roll and Vedel 1982). Additionally, such vibration techniques have been used to replay microneurographically recorded signals from muscle spindle afferents, to reproduce afferent firing patterns, enabling participants to recognise complex movement patterns (Albert et al. 2005, 2006). Such studies clearly demonstrate that these sensors are sufficient for the perception of complex and specific changes in limb posture and velocity, and do not merely elicit vague sensations. The type Ia afferents are generally described as signalling muscle stretch velocity, while the type II afferents signal absolute muscle length (Proske and Gandevia 2012), although they likely overlap substantially in their function (Macefield and Knellwolf 2018). It is, however, generally assumed that type Ia sensors are particularly important for faster movements, while type II afferents are central for estimation of maintained posture and more gradual changes (Banks et al. 2021).

This brief introduction is not exhaustive: Golgi tendon organ afferents, skin-stretch receptors and joint receptors also contribute meaningfully to proprioception. We have here focused on muscle spindle afferents in order to exemplify how position- and velocity-based sensory feedback enters the sensorimotor system.

### 2.2 Sensory integration and state estimation

Proponents of modern theories of sensorimotor control, such as Optimal Control Theory (and derived frameworks) and Active Inference, generally argue the brain assumes that sensory feed-back arises from a common hidden cause and attempts to infer its state (Wolpert et al. 1995; Todorov and Jordan 2002; Friston et al. 2010). That is, the brain integrates position-, velocity- and vision-based feedback in a self-consistent way, so that the signals ideally augment and filter one another rather than produce distinct perceptions of the same arm. A classical example of this effect in action is seen in the rubber hand illusion (Botvinick and Cohen 1998).

The presence of sensory and motor noise as well as transmission delays is fundamental to the nervous system (Van Beers et al. 2002; Faisal et al. 2008), underscoring the necessity for effective mechanisms to integrate signals across sensor types, sensory modalities and time. Bayesian sensory integration has been proposed to meet this requirement (Knill and Pouget 2004), providing a principled way to combine multiple sources of uncertain and noisy information, weighting each by its assumed state-sensor precision. Several studies in which research participants have been required to combine different perceptual cues and in which the reliability of some cues has been manipulated have provided strong support for such Bayes-optimal sensory integration (Körding and Wolpert 2004; Chancel et al. 2016; Limanowski and Friston 2020; Chancel and Ehrsson 2023). The perceived state may then further be integrated over time through learned laws of motion or generative models; for example, the arm may be expected to move smoothly through space, with congruent changes in position, velocity, and acceleration, rather than disappear in one spot and reappear at another (Friston 2008; Friston et al. 2010).

This formulation shows how velocity-based proprioceptive feedback can contribute to perceived limb position without encoding position directly. If the hand’s starting position is well represented (for example, through visual feedback), precise velocity information allows precise inference of positional change relative to that starting point.

Reaching tasks are a central tool for investigating this issue. When reaching to the position of a visual target without visual feedback of the hand, the participant implicitly indicates that perceive the hand to be at the target’s position. An endpoint error therefore indicates an offset between the perceived and actual hand position. Several studies have indicated that a currently perceived position may well be inferred relative to a previously perceived position; for example, endpoint errors are reduced when vision of the hand is provided prior to movement onset, compared to not seeing the hand at the start position (Rossetti et al. 1994; Elliott and Calvert 1990; Vindras et al. 1998). Additionally, repetitive movements remain of consistent length and direction across many repeated sequential reaches, even when absolute position errors have accumulated substantially (Brown et al. 2003a,b; Patterson et al. 2017). Similarly, position offsets induced by brief visual perturbations lead to biased endpoint errors in the opposite direction, indicating that the perceived hand position is inferred as a change from the presented visual input, even when this feedback is no longer available (Körding and Wolpert 2004; Hewitson et al. 2018).

Together, these findings suggest that estimates of hand position are updated over time using both prior state estimates and incoming sensory feedback. Bayesian state-estimation models formalize this process as a cycle of forward prediction and sensory correction (Wolpert and Ghahramani 2000). However, the relative contributions of information about absolute position and information about movement-related change to state estimation remain unclear. A brief visual offset provides a way to probe these contributions: if estimates rely strongly on movement-related information, the induced position error should persist as the estimated state is updated from the previous estimate; more precise positional feedback should correct it more rapidly. Here, we test these predictions in continuous target-tracking and sequential reaching, and compare participants’ movements with simulations that vary the assumed precision of position- and velocity-based feedback.

## 3 Methods

The study was conducted at the Department of Psychology, University of Copenhagen, and was approved by the departmental ethics review board (no. IP-EC-030723-01). Twenty-two healthy participants (16 women and six men) were enrolled in the study, all of whom were right-handed, with a mean age of 23 years (range 20–29). Written informed consent was obtained from all participants before participation in the experiment. All data were collected during a single 3-hour session, and participants were compensated at a rate of 160 DKK per hour. The data for the two tasks described here were collected as part of a larger study.

### 3.1 Experimental setup

The participant was seated at a table, the height of which was adjusted so that when the right hand rested on the table with the upper arm in a vertical position, the elbow was slightly extended past 90^*°*^. This posture helped prevent the lower arm would not slide across the tabletop during the two tasks, preventing any additional tactile feedback regarding the executed movement.

The participant was outfitted with a Meta Quest 3 virtual reality (VR) headset (Meta Platforms Technologies Ireland Limited, Dublin, Ireland), which was connected to the experiment PC via Quest Link. The experiment software was programmed in the Unity game engine (Unity Technologies, San Francisco, USA), using the Unity Experiment Framework (Brookes et al. 2020) for the trial and block structure. The VR headset was set up with a refresh rate of 90 Hz, which controlled both the refresh rate of the displays and the rate at which the system reported the tracked position of the controller. The right-hand VR controller was seated in a 3D-printed mount with a flat base with a felt slider attached underneath. This allowed the VR controller to slide across the tabletop with minimal friction, and participants were instructed to always keep the controller on the table, effectively limiting movement of the controller to a horizontal 2D plane. This was monitored in the tracking data; no trials had to be rejected due to a lifted controller. The left-hand VR controller was attached to the tabletop, which allowed table to be represented in the virtual environment. This virtual table was always visible, ensuring stable visual feedback regarding the working environment.

When visual feedback of the controller location was provided in the virtual environment, it consisted of a simple sphere (radius 1.25 cm), thereby providing position but not orientation feedback. The sphere was centred vertically on the controller’s upper surface, at the thumb button, on which the thumb was resting; this thumb button was also used as input during the two completed tasks. The visual target in both tasks was a sphere (radius 2.5 cm), positioned at the same height off the table; the centre of the controller sphere could always be aligned with that of the target sphere. Two different tasks were completed, a target-tracking task and a sequential reaching task.

#### 3.1.1 Target-tracking task

During the target-tracking task, the target moved at a constant speed along a circular path with a radius of 15 cm, completing each revolution in 4 s (see Fig. 1). The participant indicated readiness by placing the right VR controller at the nearest point on the target path, approximately 10 cm in front of their chest, and pressing the controller’s thumb button. At this point, the visual feedback of the controller would be removed, leaving only the table and the target in view. The target began at the point on the circle diametrically opposite the hand start location, 30 cm away, and moved clockwise (as viewed from above). After 2 s, it reached the start location, at which point the participant was instructed to track the target by keeping the thumb button as close to the centre of the target as possible. The target continued for 18 s (4.5 revolutions after tracking onset). A total of 40 trials were completed per participant. In three-quarters of these, visual feedback of the controller location was reinstated for 1 s, in the last quarter of the first revolution. Visual feedback was assigned pseudorandomly to be either correctly mapped or offset by 5 cm to either side.

**Fig. 1.**
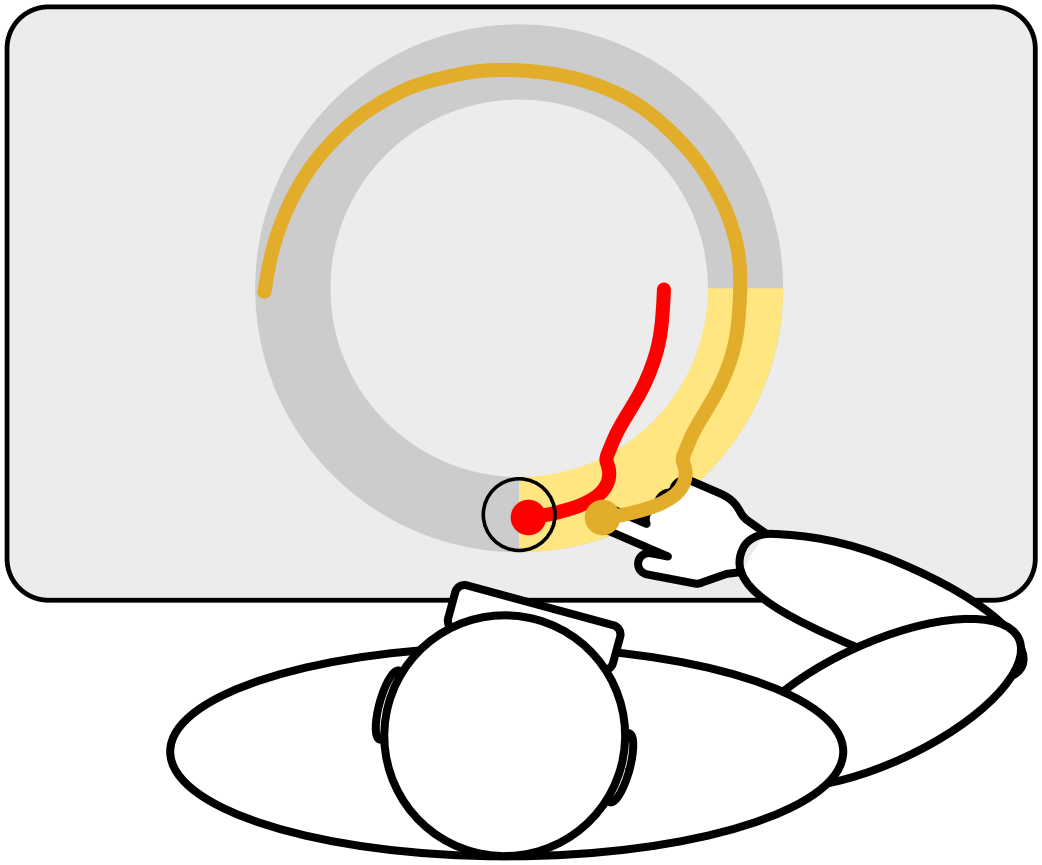
Visualisation of the target-tracking task. The research participant is seated at a table and wears a VR headset. Their task is to track a moving target (shown as a small black circle) that follows a clockwise circular track (not shown in VR, large grey circle). The target is followed for 4.5 revolutions, with each revolution lasting 4 seconds (total of 18 seconds). During the last quarter of the first revolution (shown as a yellow highlight), 1 second of visual feedback is provided, which may be offset by 5 cm to the left or right (red trace, shown here offset to the left, prompting a corrective movement to the right). The orange path shows the real movement of the tracked VR controller, which is not visible in the VR environment.

#### 3.1.2 Sequential reaching task

Because target-tracking is not a commonly used task, we also wished to approach our research question with a more standard sequential reaching task. During this task, the same start location was used, which was marked with a grey sphere, with a second grey target placed 38 cm in front of it (both with a radius of 2.5 cm). The trial was similarly initiated by pressing the thumb button when at the start location, which removed visual feedback of the controller location. After a random delay of 500–1000 ms, the far target turned green, which served as the “go” signal. From here, the participant was instructed to indicate the far target as precisely as possible by sliding the controller to it and pressing the thumb button again; this would switch the colours of the two targets, such that the start location would now be green, prompting a move to this target. This would be repeated for a total of three movements (out, back and out again). Visual feedback was reinstated for 100 ms after passing the 10 cm mark on the first outbound reach; it was either correctly mapped or offset to either side, corresponding to a 7.5^*°*^ rotation around the start position (approximately 1.3 cm lateral offset at the 10 cm mark). A total of 60 trials were completed: 20 repetitions of each condition, in pseudorandom order.

To familiarise participants with the procedure and required movement, they completed five practice trials before each task, with continuous, correctly mapped visual feedback.

### 3.2 Simulation model

We use an implementation of a computational model of a two-joint planar arm described previously (Mortensen et al. 2025). The goal of the computational model is to compare its simulated movement patterns on the above-described targettracking task and sequential reaching task with human data, while allowing us to vary the relative reliance on position- and velocity-based proprioceptive feedback. The following section provides a brief overview of the simulation approach; for a detailed description of the implementation, please refer to the cited paper or the Code Availability section (5.2).

The upper arm rotates around a fixed shoulder position and connects to the lower arm at the elbow, which flexes in only one direction. The end of the lower arm thus represents the hand position, and the entire arm is limited to movement in the horizontal plane; the diagram in Fig. 1 matches the simulated setup. The state of the arm is thus fully defined by the angular positions and velocities of the shoulder and elbow joints. While this simulated “elbow-out” configuration does not perfectly match the participants’ posture (their shoulders were well above the table), we consider it a reasonable approximation for the present purposes.

The agent controlling the simulated arm lacks access to its true state and instead receives noisy position- and velocity-based proprioceptive feed-back corresponding to the angular positions and velocities of the shoulder and elbow joints. The agent also receives visual feedback about the hand position as noisy 2D Cartesian coordinates when this is available in the simulated task. For our implementation of recursive Bayesian state estimation, we use an unscented Kalman filter (UKF) (Wan and Van Der Merwe 2000; Julier and Uhlmann 2004). In addition to noisy sensory feedback, the UKF also uses efferent motor commands generated by a linear quadratic regulator (Athans 1971). These commands are subject to signal-dependent noise (Harris and Wolpert 1998) before being used to drive the shoulder and elbow joints of the simulated arm. The simulation model is therefore conceptually similar to the model presented by Todorov and Jordan (2002) in their seminal paper on optimal feedback control. All sensory and motor noise was modelled as Gaussian white noise, with a normal distribution and no temporal autocorrelation.

A key feature of the UKF is that the relative influences of different sources of sensory feedback and efferent motor commands are weighted by their precision; a feedback channel expected to be noisier has a proportionally smaller effect on state estimation than a more precise source of information. Additionally, the current state estimate is propagated through time, thereby producing a prior against which the next set of sensory feedback is weighted. For example, the positional prior is based not only on the previous positional estimate but also on the velocity estimate, scaled by the time difference. We provide an extended example of this in Appendix A.

In our simulations, we use the same two configurations of proprioceptive noise that we presented previously in Mortensen et al. (2025). The configurations use either high-precision velocity feedback and low-precision position feedback (HVLP), or low-precision velocity feedback and high-precision position feedback (LVHP); this allows us to compare proprioceptive systems that rely primarily on velocity or positional feedback. Specifically, in the HVLP configuration, the velocity sensors have 16 times the precision of the position sensors, corresponding to a standard deviation ratio of 1:4 (velocity:position; precision = 1*/σ*^2^). This ratio is reversed in the LVHP configuration; all other settings are identical. As a consequence of Bayesian state estimation, each configuration is expected to have the following characteristics:

#### High-precision velocity, low-precision position (HVLP)

The agent estimates its velocity precisely, allowing it to infer positional changes accurately. Noisy positional feedback will have little influence on these estimates in the short term. Moment-to-moment changes in position are smooth, but small errors in velocity estimates can accumulate, and any error in the inferred position will tend to be propagated forward in time.

#### Low-precision velocity, high-precision position (LVHP)

The agent will struggle to estimate velocity and will therefore have broader, more uncertain priors for position. The comparatively precise position sensor will have a much greater effect on the posterior position estimate. Moment-to-moment changes in position will be less smooth, but any drift from the position indicated by the sensor will quickly be corrected.

We simulated the movement patterns of both of these model configurations on the same two tasks completed by the human participants. The presented results are based on 100 simulated trials per task.

### 3.3 Statistics

We quantified the time course of the visually induced offset in the target-tracking task by fitting a hierarchical exponential-decay model using the brms package (Bürkner 2017) in R (R Core Team 2024). We used a nonlinear Gaussian model:

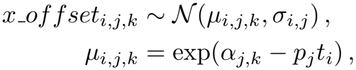

where *x_offset*_*i,j,k*_ is the *x*-axis offset for observation *i*, participant *j*, and visual-offset condition *k*; *t*_*i*_ is time in seconds since visual feedback was removed; *α*_*j,k*_ is the log-amplitude at *t* = 0; *p*_*j*_ is the participant-specific decay rate; and *σ*_*i,j*_ is the residual standard deviation for participant *j*. Thus, the expected offset at *t* = 0 is exp(*α*_*j,k*_), and larger positive values of *p*_*j*_ correspond to faster decay. The implied half-life for participant *j* is log(2)*/p*_*j*_.

Before fitting the model, we calculated the offset along the *x*-axis for each trial by subtracting, at each time point, the participant’s mean x-position in the correctly mapped visual-feedback condition from the x-position in the offset condition. This within-participant centring accounts for any consistent error in the movement path, relative to the target. We sign-aligned the two offset directions by multiplying offsets from one direction by *−*1, so that positive values represent the induced offset in both conditions. We fitted a shared decay rate distribution of to the two sign-aligned conditions, while allowing their initial amplitudes to differ.

The condition-specific log-amplitudes were modelled as

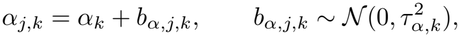

where *α*_*k*_ is the population-level log-amplitude for condition *k*, and *τ*_*α,k*_ describes between-participant variation in that condition. The decay rates were modelled as

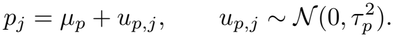

The residual standard deviation was also allowed to vary between participants on the log scale.

Priors on all population-level parameters and standard deviations provided weak regularisation; see the Code Availability section (5.2) for details. The model converged without divergent transitions; all 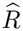 values were below 1.005, and visual inspection of caterpillar plots indicated no fitting issues.

## 4 Results

### 4.1 Target-tracking task

Fig. 2**a** and **b** show the simulated movements of the Bayesian agent for the LVHP and HVLP configurations, respectively, while **c** shows the corresponding human data, all from the target-tracking task. The light grey circle in each subplot indicates the path taken by the tracked target, which always moved clockwise, with each full revolution lasting 4 s. The three coloured traces represent the averaged movement paths: the baseline condition with correctly mapped visual feedback (green), and the two offset conditions (5 cm to the right or left), which prompted corrective movements in the opposite direction (blue and orange, respectively). The column covering 3–4 s shows the 1 s period during which visual feedback of the hand position was available, including the corrective movements performed by human participants and both agent configurations.

**Fig. 2.**
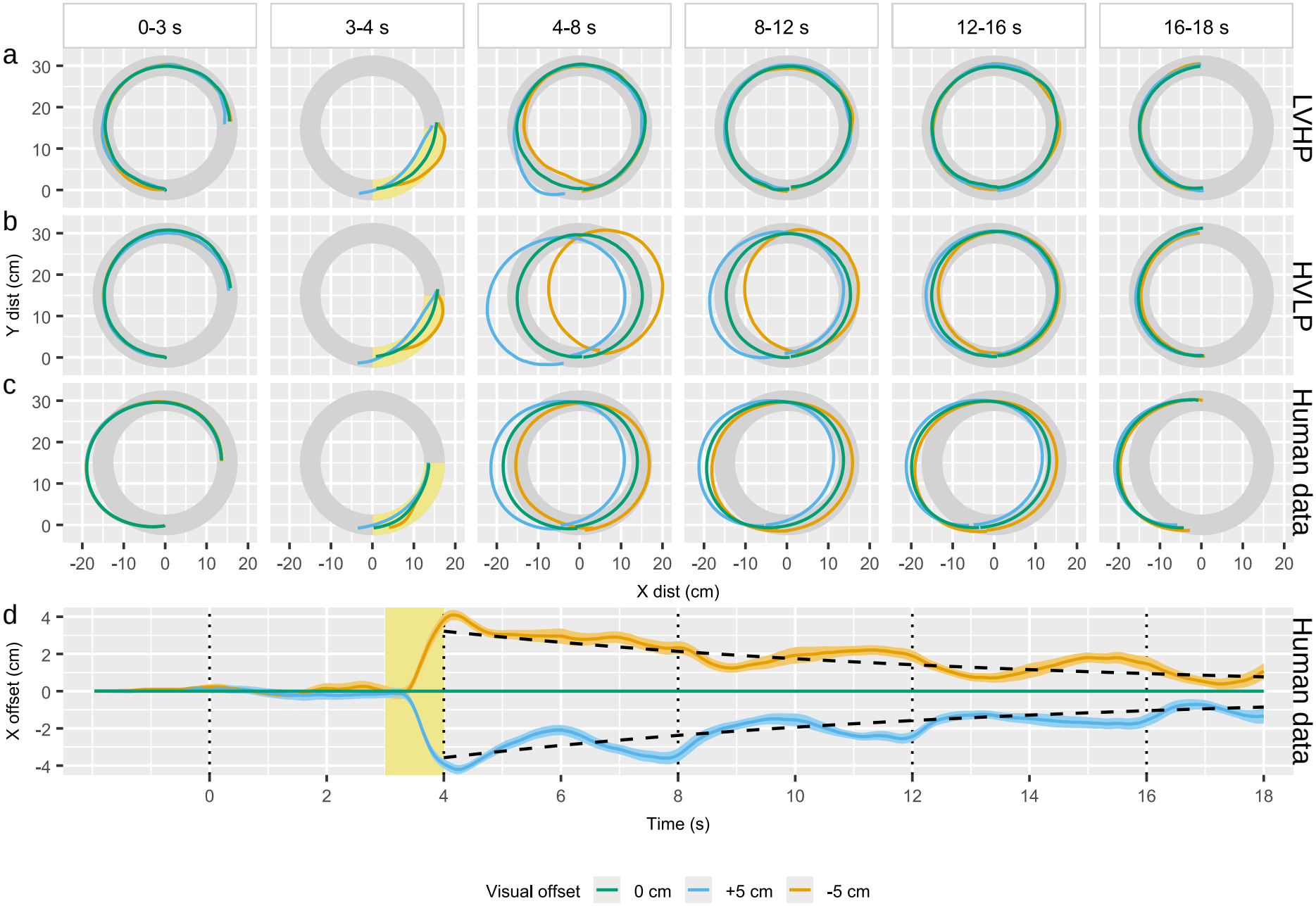
Results of simulated and human participants in the target-tracking task. Subplot rows **a** and **b** show the mean movement paths of the LVHP and HVLP proprioceptive configurations, respectively, each averaged across 100 simulated trials, while subplot row **c** shows the corresponding group mean movement paths of the human participants. The grey circle in each subplot shows the path of the moving target, while the three coloured traces show the correctly mapped condition and the two visual-offset conditions, as indicated in the legend. Movement direction was always clockwise, and visual feedback was provided only during the 3–4 s interval (yellow highlight). Panel **d** plots the mean induced offset in the movement path along the *x*-axis for the human participants, calculated by subtracting the mean *x*-position in the control 0 cm-offset condition from the mean *x*-positions in the other two conditions, within participant. The shaded bands show the group mean and *±*1 standard error of the mean (SEM) across participants, calculated from participant-level mean offsets at each time point. The dashed black lines indicate the fit of the group-mean exponential-decay model, while the vertical dotted lines, spaced every 4 s, show when the tracking target passes the 6-o’clock start position.

Focusing first on the human participants, the subplots covering the 4–18-second interval in Fig. 2**c** show that the tracking movement tends to remain offset after visual feedback is removed, with the gap between the offset and non-offset conditions slowly decreasing over the following revolutions. This is further highlighted in Fig. 2**d**, which plots the slow decay of the induced positional offset along the *x*-axis relative to the baseline condition.

To quantify the time dynamics of this process, as well as the between-participant variability, we fitted a hierarchical exponential decay model, as described in Section 3.3. The model revealed a group-mean decay rate (*p*) of 0.10 (95% credible interval (CrI) [0.08, 0.12]), corresponding to a half-life of the induced offset of 6.9 seconds (95% CrI [5.8, 8.7]). The between-participant variability in the decay rate was estimated as a standard deviation (SD) of 0.05 (95% CrI [0.04, 0.07]), with posterior individual *p* estimates ranging from 0.02 to 0.21. These correspond to estimated half-lives of 3.3 and 40.6 seconds for participants with the fastest and slowest estimated correction rates, respectively. The fit of the group-mean exponential decay function is plotted as a dashed line in Fig. 2**d**.

The persistence of the induced offset during tracking after visual feedback was removed indicates that perceived hand position is updated in a way that depends not only on the latest sensory feedback but also on previously estimated states. Additionally, it appears that velocity-based information contributes substantially to this process; relative movements remain well-formed, with the hand continuing along a circular path that is slightly offset from that of the baseline condition. If absolute positional proprioceptive feedback were both highly precise and heavily weighted for state inference, we would expect the visually induced offset to be discarded much more quickly after visual feedback was removed. The tendency of the movement paths of the three tested conditions to converge back towards each other, however, suggests that absolute positional information is available for sensory integration. The slowness of this convergence may be interpreted as reflecting the relatively low weight assigned to positional feedback during sensory integration.

Compared with the two simulated agent configurations, the human data show a pattern similar to that produced by the HVLP configuration. We cannot be certain that the explanation proposed above accounts for the pattern in the human data, such as we can in the case of the simulated data. As a counter example, when relatively high-precision positional feedback is available in the LVHP configuration, the induced offset in the agent’s posterior state estimate diminishes sub-stantially within 1–2 seconds after visual feedback is removed.

Both simulated agents estimate that they are following the target accurately throughout each trial. Corrections in response to offset visual feedback are completed within the 3–4 s window; thereafter, both detect minimal discrepancies between their hand position and the target, irrespective of the visual condition (not shown).

The offset plotted in Fig. 2**d** also shows an oscillating pattern with a period of approximately 4 seconds. This occurs because the induced offset is not a constant Cartesian offset throughout the circular movement. Although we cannot confirm the exact reason for this effect in our human participants, a similar effect appears in the simulated data, most clearly in the 4–8 s column of Fig. 2**b**, where the blue trace is skewed down and to the left while the orange trace is skewed up and to the right. This occurs because state estimation is performed in the agent’s proprioceptive reference frame, and a particular angular mis-estimation in the joints does not produce a constant Cartesian offset across the arm configurations used during the task.

#### 4.1.1 Example individuals

To highlight variability in individual responses, we show the mean movement paths of two participants in Fig. 3. We selected two participants with contrasting movement patterns and responses to the visual offset. Example **a** in the top row of Fig. 3 shows a strong tendency to move relative to the visually induced offset, with clear differences between the three visual conditions remaining until the end of the tracking movement.

**Fig. 3.**
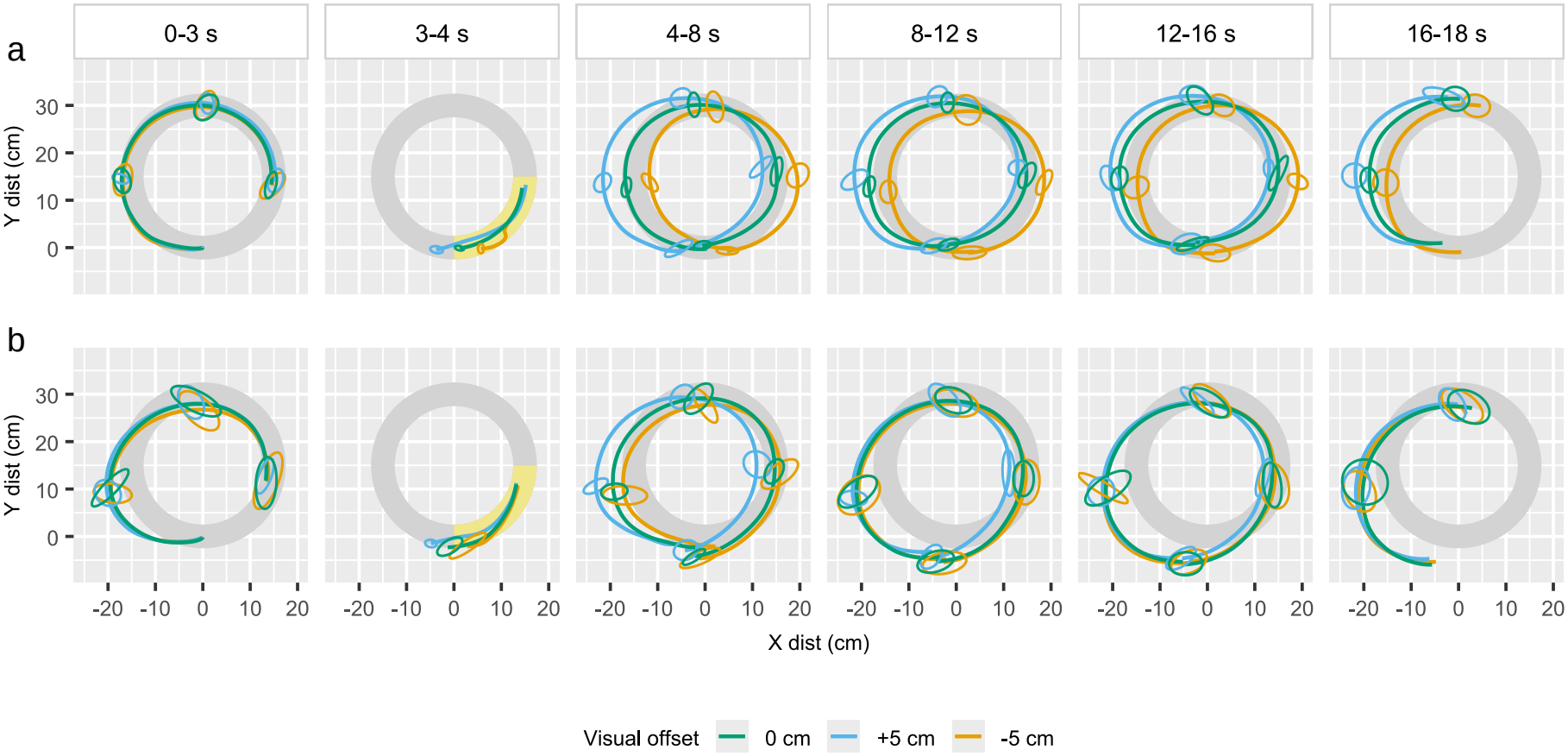
Mean movement paths for two example participants in the target-tracking task. At one-second intervals (four times per revolution), ellipses show the within-participant variability in hand position across trials. Each ellipse represents the 1-SD contour of hand positions at that time point. Appendix B shows the corresponding plots for the remaining 20 participants.

Example **b**, on the other hand, appears to be minimally affected by the visual offset just 4 seconds after visual feedback is removed. This example also illustrates how some participants consistently deviate from the shape of the circular movement path, potentially indicating distortions and/or constant offsets between their proprioceptive and visual reference frames. The plotted ellipses showing within-participant variability in movement show the consistency of this type of error. Similarly, many other participants tended to follow a circle that was too small or too large, or a distorted circle; see Appendix B, where the data for the remaining 20 participants are plotted.

#### 4.1.2 Perceptual drift

Although the offset conditions gradually converge towards the non-offset condition along the *x*-axis, this provides only a simplified description of the overall drift apparent in our data. In Fig. 4, we plot how the hand position changes over the course of the target-tracking task as the target passes the same position multiple times. To illustrate this, we zoom in on the left edge of the target’s path and plot the group-mean hand position when the target passes the nine o’clock position on each revolution. The points represent 5, 9, 13 and 17 seconds after movement onset, respectively. For the three conditions with visual feedback (green, blue and orange traces), the first point is 1 second after feedback was removed. We also plot the condition with no visual feedback (purple trace).

**Fig. 4.**
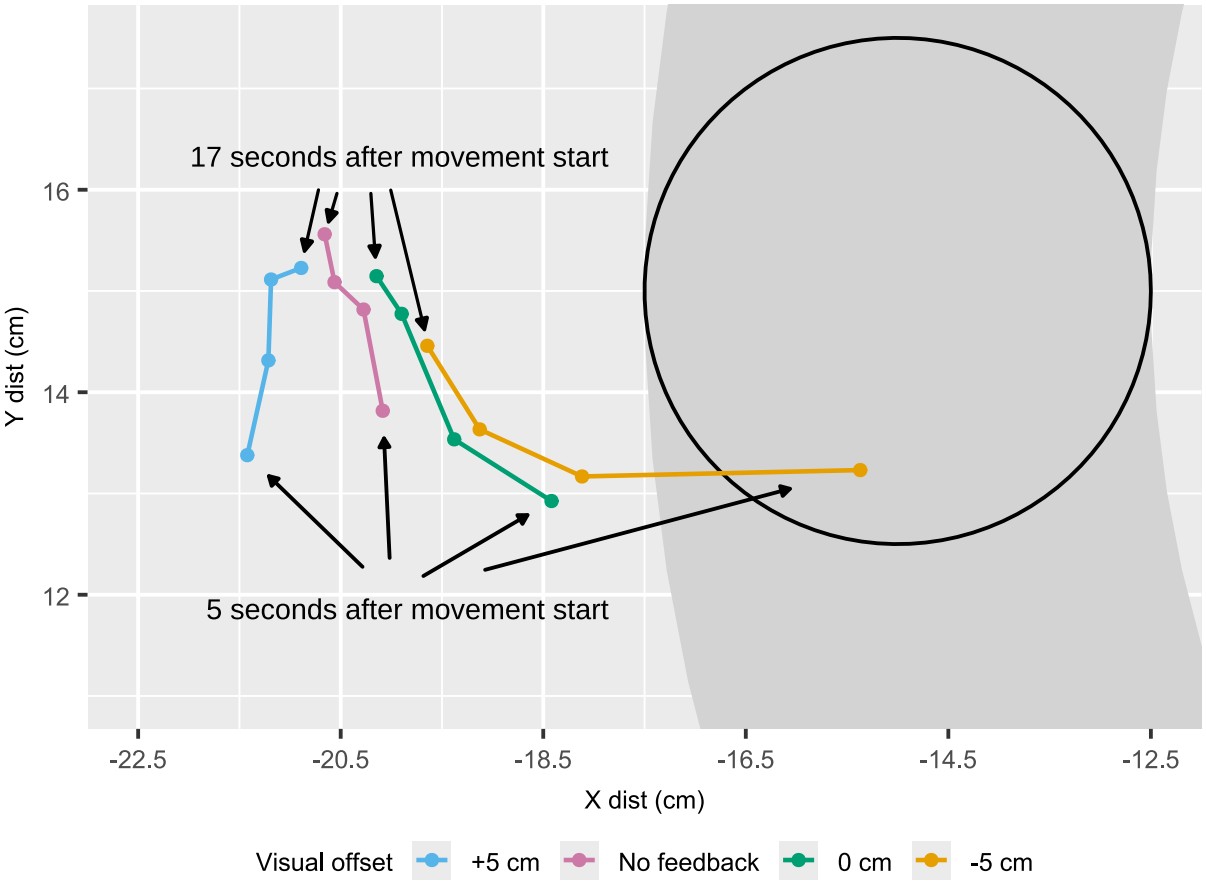
Perceptual drift during the target-tracking task. The figure illustrates positional drift during continuous target tracking, measured each time the target passes the same position over the course of a trial. Each trace shows the change in group-mean hand position over the course of the target-tracking task for each of the four visual-feedback conditions. All points represent the hand position when the target passes the nine o’clock position on each revolution. The points represent 5, 9, 13 and 17 seconds after movement onset, respectively; the first is 1 second after visual feedback is removed (excepting the purple condition where visual feedback was not reinstated). The outline of the target and its path is plotted for scale.

This illustrates how the perceived hand position drifts over the course of the tracking movement, with all four conditions converging towards a single point, showing the same pattern as Fig. 2**d**.

### 4.2 Sequential reaching task

Fig. 5 shows the results of the sequential reaching task completed by the human participants (**a**), as well as the simulated movements produced by the two agent configurations (**b** and **c**). Visual feedback was withheld, except for a 100 ms interval after participants passed the 10 cm mark on the first outward reach; it was either correctly mapped or rotated by 7.5^*°*^ to either side around the starting point (an approximate lateral offset of 1.3 cm at the point where feedback was shown). The period with visual feedback is indicated by the yellow highlight. However, because participants were instructed to move at their own speed and focus on indicating the target as precisely as possible, the spatial location at which visual feedback was removed varied across participants and trials. The offset visual feedback is also shown as bold lines in the corresponding colour; for example, the visual offset to the left is plotted in orange, prompting a corrective movement to the right.

**Fig. 5.**
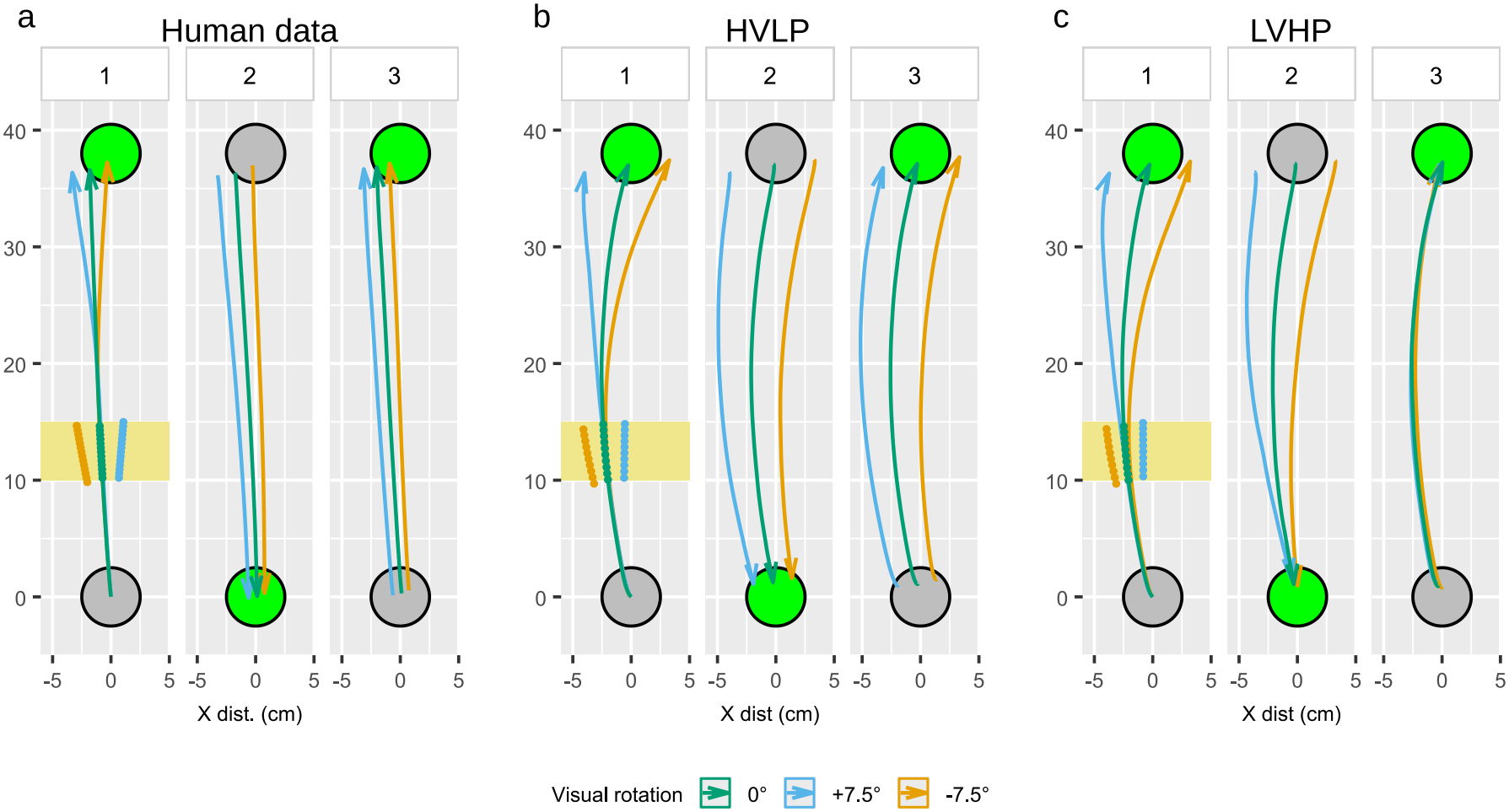
Results of simulated and human participants in the sequential reaching task. The figure shows the mean movement paths across the three sequential reaches towards targets placed 38 cm apart. Visual feedback is withheld, except for 100 ms after passing the 10 cm mark during the first reach, illustrated with the yellow highlight. Visual feedback was rotated by either 0^*°*^, +7.5^*°*^ or *−*7.5^*°*^ around the starting position; the offset visual feedback is shown in the yellow highlight. Subplot **a** shows the group mean movement path of the human participants, while **b** and **c** show the simulated movement paths from the HVLP and LVHP agent configurations, respectively. Each subplot contains three panels, one for each movement in the out-back-out sequential reaching task.

The group-mean movement paths of the human participants plotted in panel **a** show much the same pattern as that seen in the circular tracking task: the visually induced offset continues to affect the movement path during subsequent reaches.

The HVLP and LVHP configurations both produce similar movement paths during the first outward reach, both showing a style of correction similar to that of the human participants, but differ substantially over the next two movements. The HVLP configuration retains the induced offset across all three movements, as do the human participants, whereas the LVHP configuration is no longer affected by the visual feedback by the time the second target is reached. The task was simulated with 1 s per reach, corresponding approximately to the group mean time per reach of 0.99 s; individual means ranged from 0.48 s to 1.65 s.

## 5 Discussion

When reaching to or tracking a visual target without visual feedback of the hand, participants’ movements provide an indirect measure of how they estimate hand position relative to the target. We used this approach to test how visually induced position offsets persist after visual feedback is removed. Under Bayesian sensory integration, precise positional feedback should rapidly correct an offset, whereas greater reliance on rate-of-change signals should allow the offset to persist by carrying forward the previous state estimate.

The central finding of the current study is that estimates of hand position and arm state appear to be history-dependent: a visually induced offset during a motor task continues to affect the executed movement path long after the perturbing stimulus is removed. In the target-tracking task, the group-mean decay half-life was 6.9 seconds, with substantial individual variation. This exponential fit provides a compact description of the decay of the offset, rather than a direct estimate of the underlying Bayesian filtering process: positional correction depends not only on the relative precision of positional and velocity signals but also on the changing uncertainty of the current state estimate. Thus, the rate of correction may vary over the course of a movement.

These results are consistent with Bayesian frameworks of state estimation, which are central to both Active Inference (Friston et al. 2010) and Optimal Feedback Control (Todorov and Jordan 2002), in which the current state estimate is passed forward in time to act as a prior for new sensory feedback. Our modelling results further build on these widely used frameworks by highlighting the need to consider carefully the composition of proprioceptive feedback. Proprioceptive feedback is often implemented as purely positional, with time-derivative estimates computed downstream from the sensory signals (e.g. Friston et al. 2010; Limanowski and Friston 2020; Maselli et al. 2022). While the above-cited work has elegantly demonstrated how Active Inference can accommodate position-only feedback in a variety of setups, our simulation results highlight the types of tasks and situations in which the distinction between absolute and rate-of-change-based feedback may matter for inference, and offer a way to relate these frameworks to the specialised roles of type II and type Ia muscle spindle afferents.

Bringing our experimental and simulated results together, human behaviour appears consistent with a model of Bayesian sensory integration that relies heavily on velocity-based proprioceptive feedback while receiving comparatively imprecise positional feedback. As we have previously demonstrated (Mortensen et al. 2025), this model is also able to account for the reaching corrections and endpoint errors reported by Körding and Wolpert (2004). In the present work, we extend this type of reaching task to show that perceptual offsets induced by biased visual feed-back lasting only 100 ms persist not only during the ongoing movement but also across several sequential movements. This interpretation is also consistent with studies showing that visual feed-back of the hand before movement onset improves endpoint precision (Rossetti et al. 1994; Elliott and Madalena 1987; Vindras et al. 1998). This suggests that endpoint estimates depend partly on how accurately the starting position is represented, alongside information about the movement that follows.

However, although our model’s ability to reproduce human behaviour shows that its assumptions can reproduce the observed patterns in these tasks, it does not establish that humans use the same mechanism.

### 5.1 Perceptual drift and correction

Proprioceptive drift has been observed during repeated movements without visual feedback of the hand: absolute hand position can shift over time even while successive reaches remain relatively consistent (Brown et al. 2003b,a; Patterson et al. 2017). Our findings suggest a close relationship between this drift and the gradual correction observed in our results. However, the approaches differ: prior work documented drift emerging without a specific imposed positional discrepancy, whereas our brief visual perturbation induced a controlled offset whose persistence and decay relative to each other could be tracked. In both cases, movement may be guided by updating a prior estimate of hand position with movement-related information, while positional feedback gradually corrects accumulated error.

Findings from Smeets et al.’s sequential-reaching study further suggest that drift can be stable within individuals and may reflect a persistent mismatch between visual and proprioceptive reference frames, rather than random sensory or movement error alone (Smeets et al. 2006). This offers one possible context for the gradual realignment and the consistent within-participant errors that we observe, although our experiment does not establish that the same source produces both effects.

Our simulations show how this pattern can arise when movement-related signals strongly update position estimates while positional feedback corrects them more slowly. The high-precision-velocity, low-precision-position configuration reproduces the persistence and gradual correction of the induced offset. However, because the model assumes an ideal mapping between its state and sensory observations, it does not reproduce drift toward a stable baseline offset. Future work could model systematic mismatches between visual and proprioceptive reference frames.

### 5.2 Sources of movement-related information

Type Ia and type II muscle spindle afferents are plausible sources of movement-related and position-related proprioceptive information, respectively (Proske and Gandevia 2012; Banks et al. 2021). However, our task cannot isolate their contributions from information about movement carried by efferent motor commands. Both sources could support the forward updating of position estimates, so our results do not identify their relative contributions. Consistent with this ambiguity, in our model, increasing the precision assigned to motor-command information while decreasing that assigned to velocity-based feedback, or vice versa, could produce similar movement patterns. Evidence that antagonist muscle vibration can bias perceived movement and alter reaching kinematics supports a role for Ia-related signals (Cody et al. 1990; Eschelmuller et al. 2023; Mortensen and Christensen 2025; Proske and Gandevia 2012), but does not establish the contribution of Ia-related signals relative to motor commands.

Together, these findings show that visually induced hand-position offsets can persist during continuous tracking and sequential reaching, and that a Bayesian model with strong movementrelated updating and less precise positional feedback can reproduce this pattern. The results support history-dependent state estimation, but do not identify the relative contributions of proprioceptive and motor-command signals.

## Acknowledgements

This project was supported by a DATA+ grant from the University of Copenhagen. ESM and MSC were supported by the Carlsberg Foundation (CF22-0941). MSC was further supported by Independent Research Fund Denmark (DFFFKK, 0132-00141B). A subset of the data was previously presented as a poster at the Cognitive Computational Neuroscience conference in Amsterdam in August 2025.

## Statements and Declarations

### Supplementary information

Not applicable.

## Ethics approval and consent to participate

The study was approved by the ethics review board of the Department of Psychology, University of Copenhagen (no. IP-EC-030723-01). All participants provided written informed consent before taking part in the study.

## Consent for publication

All authors consent to the publication of this manuscript.

## Data availability

Experimental and simulated datasets are available at https://osf.io/xz5jt.

## Materials availability

Not applicable.

## Code availability

The model code is available at https://github.com/CoInAct-group/BayesianInferenceArm and archived at https://osf.io/kvtcu.

## Author contributions

ESM and MSC both conceptualised the work. ESM collected and analysed the data and drafted the manuscript. ESM and MSC both revised and approved the final manuscript.

## Appendix A Illustrating movement-based state prediction

The simulation uses an unscented Kalman filter to estimate the state of a two-joint arm under nonlinear dynamics; its full implementation is described in (Mortensen et al. 2025). Rather than present the full filter here, we use a simplified one-dimensional example to illustrate how movement-related information can carry a position estimate forward over time.

Suppose an agent estimates its position *x* and velocity 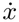 from noisy sensory observations. At each time step, it first predicts its current state from the previous estimate and the expected effects of its motor command. In a simplified model, the predicted velocity and position are

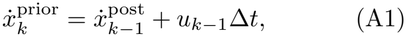

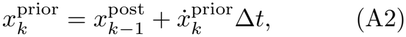

where *u*_*k−*1_ represents the expected change in velocity due to the motor command. The predicted position therefore depends on both the previous position estimate and the estimated movement since that previous estimate. If the starting position estimate is offset, accurate velocity information can preserve that offset as the agent predicts its position at the next time step.

The agent then corrects its prediction using new sensory observations. The balance between prediction and observation depends on their relative uncertainties: more precise information has greater influence on the updated estimate. Thus, precise position feedback can quickly correct an offset, whereas less precise position feedback allows prediction based on movement to carry the offset forward for longer. The offset may nevertheless diminish gradually as positional evidence accumulates.

## Appendix B Target-tracking task: Individual results

**Fig. B1.**
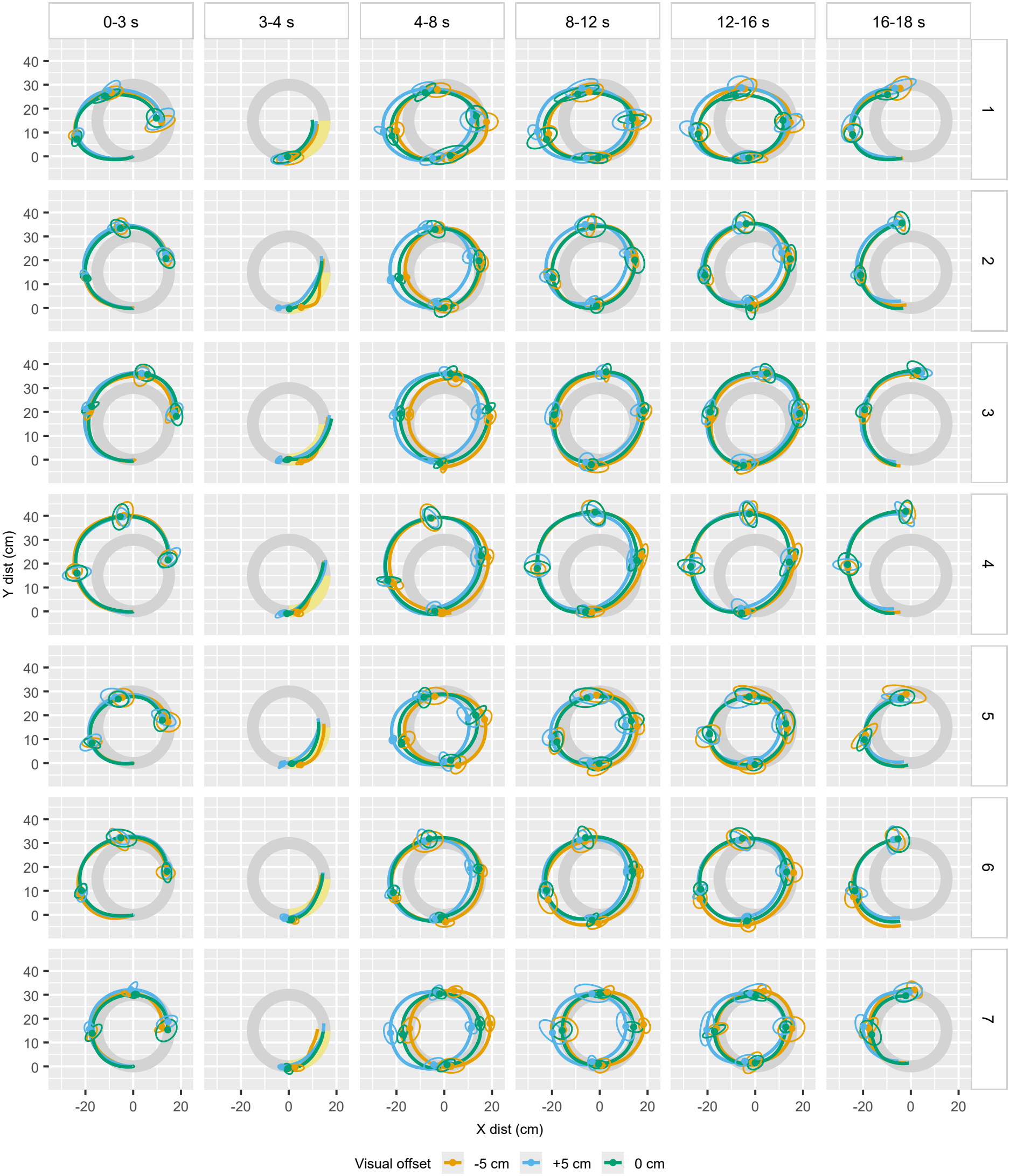
Mean movement paths for participants 1–7 in the target-tracking task. Ellipses show the within-participant variability in hand position across trials at one-second intervals. Each ellipse represents the 1-SD contour of hand positions at that time point.

**Fig. B2.**
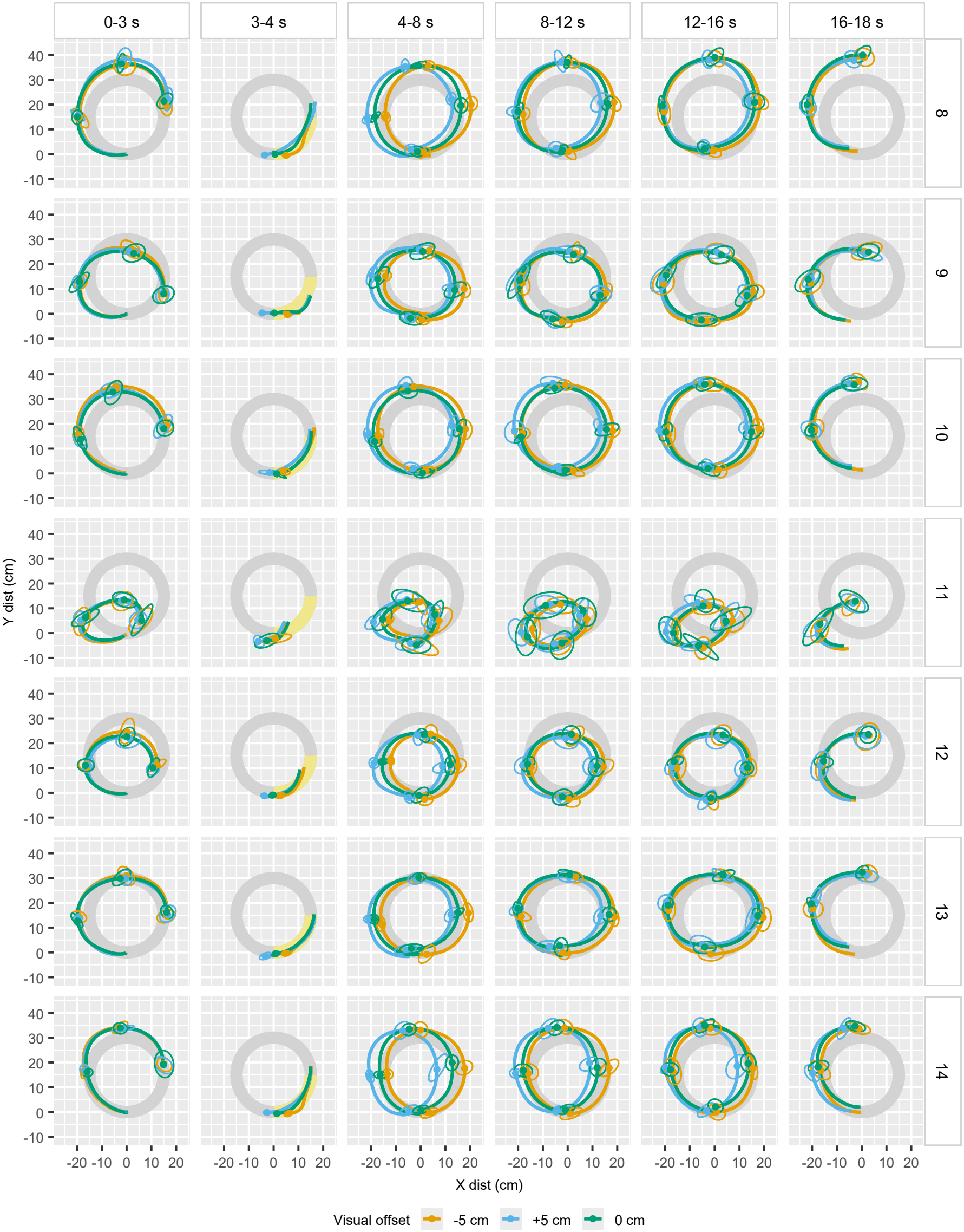
Mean movement paths for participants 8–14 in the target-tracking task. Ellipses show the within-participant variability in hand position across trials at one-second intervals. Each ellipse represents the 1-SD contour of hand positions at that time point.

**Fig. B3.**
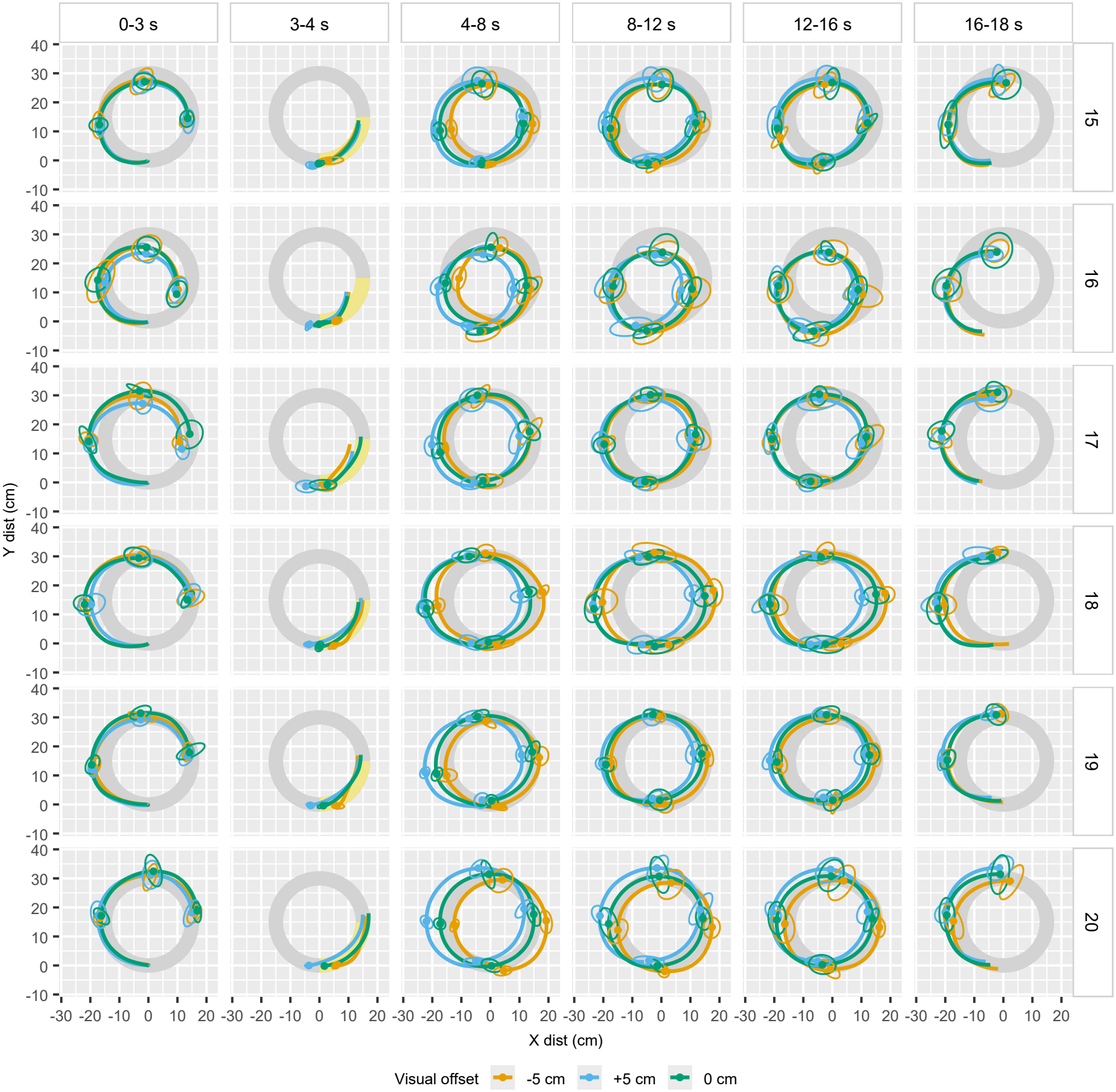
Mean movement paths for participants 15–20 in the target-tracking task. Ellipses show the within-participant variability in hand position across trials at one-second intervals. Each ellipse represents the 1-SD contour of hand positions at that time point.

